# Combined analysis of transposable elements and structural variation in maize genomes reveals genome contraction outpaces expansion

**DOI:** 10.1101/2023.03.02.530873

**Authors:** Manisha Munasinghe, Andrew Read, Michelle C. Stitzer, Baoxing Song, Claire Menard, Kristy Yubo Ma, Yaniv Brandvain, Candice N. Hirsch, Nathan Springer

## Abstract

**Background:** Structural differences between genomes are a major source of genetic variation that contributes to phenotypic differences. Transposable elements, mobile genetic sequences capable of increasing their copy number and propagating themselves within genomes, can generate structural variation. However, their repetitive nature makes it difficult to characterize fine-scale differences in their presence at specific positions, limiting our understanding of their impact on genome variation. Domesticated maize is a particularly good system for exploring the impact of transposable element proliferation as over 70% of the genome is annotated as transposable elements. High-quality transposable element annotations were recently generated for *de-novo* genome assemblies of 26 diverse inbred maize lines.

**Results:** We generated base-pair resolved pairwise alignments between the B73 maize reference genome and the remaining 25 inbred maize line assemblies. From this data, we classified transposable elements as either shared or polymorphic in a given pairwise comparison. Our analysis uncovered substantial structural variation between lines, representing both putative insertion and deletion events. Putative insertions in SNP depleted regions, which represent recently diverged identity by state blocks, suggest some TE families may still be active. However, our analysis reveals that, genome-wide, deletions of transposable elements account for more structural variation than insertions. These deletions are often large structural variants containing multiple transposable elements.

**Conclusions:** Combined, our results highlight how transposable elements contribute to structural variation and demonstrate that deletion events are a major contributor to genomic differences.

## Background

Plant genomes are replete with transposable elements (TEs) — accounting for as little as 20% of the genomes of *Medicago truncatula* and *Arabidopsis thaliana* (1) to over 70% of the genomes of maize and wheat (1–3). TE expansion is mediated by the active movement of TEs, particularly class I ‘copy-and-paste’ elements that utilize an RNA intermediate, and can contribute to expansions in genome size. Many closely related plant species have similar gene content but substantial differences in genome size attributable to TE accumulation in one species (4–7). The continued movement of TEs in many plant lineages has been hypothesized to lead to a ‘one-way ticket to genome obesity’ (8). However, this hypothesis ignores the potential of deletions to reduce TE content and genome size. In fact, on-going deletion events in many plant genomes could counteract genome size expansion caused by TE accumulation (8–13). Thus, the contribution of TEs to an organism’s genome size depends on both the frequency of TE insertions and deletions.

Advancements in whole-genome long-read sequencing and computation methods have rapidly enhanced our ability to characterize and investigate structural variation between genomes (14–16). Genome-wide characterization of structural variation in the maize genome has found extensive variability in genome sequence (17–23), gene content (22–25), and transposable element content (23,26). Detailed characterization of multiple haplotypes for several loci in maize revealed extensive structural polymorphisms for TE content (27–29). Given the high TE content of the maize genome (2), it is likely that transposable elements are a major contributor to structural variants (SVs), but this has yet to be fully quantified.

The recently completed high-quality genome sequences of the 26 maize inbred lines used to generate the nested association mapping (NAM) population provides an opportunity to generate a high-resolution understanding of transposable element polymorphisms and the extent to which variation in TEs contributes to SVs and phenotypic variation in maize (23). These lines were selected from a larger association panel to provide a sampling of maize diversity. As such, these lines have limited genetic relatedness or population structure. We generated base-pair resolved whole genome alignments in a pairwise fashion by aligning the B73 reference genome to each of the other NAM founder lines using AnchorWave (30). We developed an approach that intersected these pairwise alignments with robust, consistent TE annotations generated for each of the NAM lines (23,31). This allowed us to classify each TE annotation as either shared, polymorphic, or ambiguous between two genomes. Applying this approach revealed that >30% of TEs are polymorphic in comparisons between B73 and a given NAM genome. A comparison of all structural variants and TE annotations revealed that TEs contribute substantially to structural variation among NAM genomes, but that only a subset of the structural variants have features that suggest simple TE insertion polymorphisms. A careful examination of large genomic regions that are likely recently diverged across these comparisons identified a subset of TE families that may have ongoing movement in modern maize inbreds.

## Results

Our first goal is to quantify the number of TEs in maize genomes and how these TEs contribute to genome size. To do so, we make use of previous annotations generated using the panEDTA approach (23,31,32), which provide a set of structural annotations representing putative full-length transposons with intact structural features (long terminal repeats, terminal inverted repeats, target site duplications, etc.) as well as homology based annotations of transposon-associated sequences that contain sequence similarity to structurally annotated elements. After numerous standard quality control steps (e.g., excluding helitron annotations, non-TE repeats, and potentially misannotated TEs (see methods for details)), we found that, on average, NAM lines had 858,902 transposons. 7.9% of these transposons were structurally annotated and the remaining 92.1% relied on homology-based annotations (Table S1). The average total of transposon sequence in the NAM genomes is 1,655Mb (min=1,634Mb, max=1,673Mb) with structurally annotated TEs account for an average of 31.2% of the total TE Mb (Table S1). In contrast, only 61Mb of the NAM genomes is annotated as genes. Structural annotations account for 8.7% of all class I elements and 6.3% of all class II elements. However, 32.8% of the Mb of class I elements are structurally annotated, while only 14.1% of the Mb of class II elements are structurally annotated.

There are many difficulties in accurately annotating transposons that can complicate the exact quantification of transposon variation. For example, homology annotations or nested insertions can result in a single transposon being represented by multiple annotation fragments that are not clearly associated with a single element. This can lead to the number of transposon annotations over-representing the total number of actual transposons. This is particularly noted for longer LTR elements. While the true number of shared and polymorphic transposons can vary based on annotation quality and approaches, the cumulative base pairs of transposons that are shared or polymorphic are less subject to influences based on annotating fragments of a transposon. Consequently, we report both the number and cumulative Mb of transposon-associated sequences.

### Highly variable TE content among NAM genomes

We next aimed to characterize variation in TE content across maize lines. Pairwise whole-genome alignments generated using AnchorWave were used to classify TEs as shared or polymorphic between B73 and a singular NAM genome (30). In each pairwise contrast between B73 and a NAM genome, each region of the alignment could be classified into one of three types: alignable sequence, structural variant sequence present in one genotype relative to the other genotype, or unalignable sequence (see methods for details) (Figure 1A).

**Figure 1.**
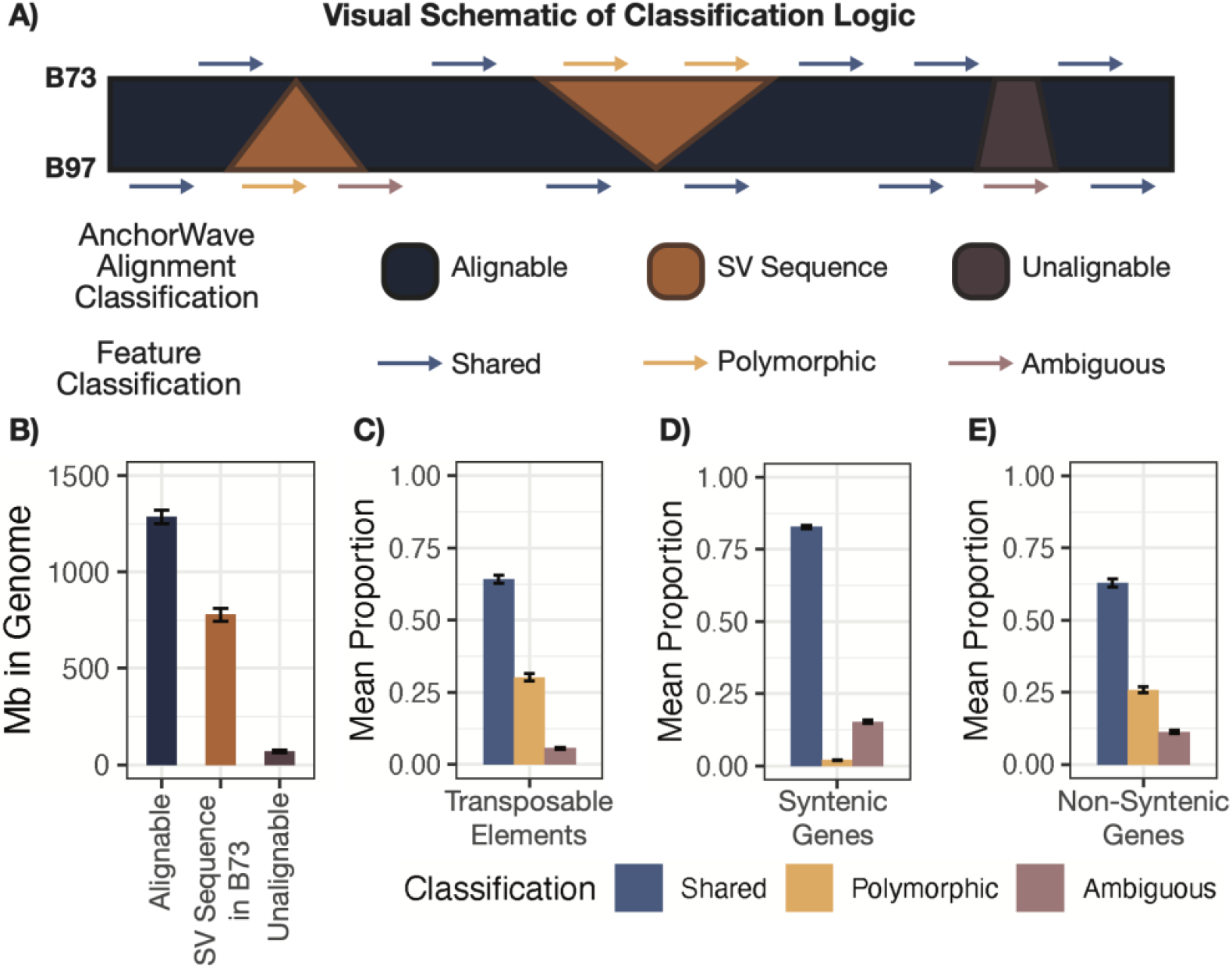
Proportions of Various Classifications Across B73 versus NAM comparisons. (A) A visual schematic describing the logic flow used to both partition the pairwise AnchorWave alignments between B73 and every other NAM line into alignable, structural variant, or unalignable sequence and to intersect these regions with feature annotations to classify features as either shared, polymorphic, or ambiguous. (B) Barplots showing the Mb of sequence in B73 classified as alignable (dark blue), structural variant sequence in B73 (orange), and unalignable (purple) across all pairwise comparison between B73 and every other NAM genome. Barplots showing the mean proportion of (C) transposable elements, (D) syntenic genes, and (E) non-syntenic genes in B73 classified as either shared (light blue), polymorphic (yellow), or ambiguous (light purple) across all pairwise comparisons between B73 and every other NAM genome. Height of the bar in panels B – E indicate the mean with error bars indicating the standard deviation across all 25 pairwise comparisons.

Across all pairwise comparisons between B73 and all other NAM lines, 60.3% of the B73 genome is alignable, 36.5% is inserted sequence in B73, and 3.2% is unalignable when compared to the other genomes (Figure 1B,S1). Reassuringly, on average, 36.1% of a NAM’s genome contains inserted sequences absent in B73 (Figure S1).

To classify TEs, the coordinates of annotated TEs within NAM genomes were compared to the AnchorWave classifications of the genome. Any TE that had ≥ 95% overlap with alignable regions was classified as ‘shared’ between B73 and the focal NAM line, while TEs that had ≥ 95% overlap with inserted sequence were classified as ‘polymorphic’ between B73 and the focal NAM line. The remaining TE annotations were classified as ‘ambiguous’. These ambiguous features include examples that are within unalignable regions as well as those that partially overlap (< 95% overlap) regions with alignable or inserted sequence. On average, 64.2% of B73 TEs were classified as shared, 30.2% were classified as polymorphic, and the remaining 5.6% were classified as ambiguous (Figure 1C). This means that, in any pairwise comparison between B73 and another NAM genome, approximately 259,000 B73 TE annotations are polymorphic or absent in the NAM genome. There is relatively little variation in the number of TEs that are classified as either shared, polymorphic, or ambiguous depending on whether we are characterizing the TEs present in B73 or the compared NAM genome, and the proportions are quite similar for each of the NAM genomes (Figure S2).

Several prior studies have evaluated the frequency of present-absent gene sequences among maize genomes using a variety of approaches (18,20,22–24). To evaluate the frequency of shared and polymorphic classifications for TEs relative to genes in a consistent fashion, we applied the same approach described for TEs above to gene annotations in maize (Figure 1D,E). The maize gene annotations were split into syntenic and non-syntenic (based on comparisons to other grasses). Non-syntenic genes often include pseudogenes or transposed gene fragments, and they are much more variable between genomes (33). As expected, we found that syntenic genes are much more likely to be shared between genomes with a relatively low polymorphic rate (Figure 1D). Non-syntenic genes, in contrast, have higher rates of polymorphic genes that are nearly as high as the rate of polymorphic TEs (Figure 1E). The frequencies of polymorphic genes based on our approach is similar to previous estimates. In general, genes exhibit a higher frequency of ambiguous classifications in comparison to TEs, but many of these likely reflect insertion/deletion events within introns that result in less than 95% of the gene sequence being present within alignable sequence. When we classify genes based only on exon sequence, we find that the proportion of ambiguous classifications is reduced (Figure S2).

The B73 TEs were compared to each of the other NAM genomes in pairwise comparisons. An analysis across each of these pairwise comparisons provides the opportunity to characterize the classification frequency for each of the B73 TEs (Figure 2A). The B73 features, either TEs or genes, could be classified as core (classified as shared across all 25 comparisons), near-core (classified as shared in 23 – 24 comparisons), variable (shared in 1 – 22 comparisons), or as private (never classified as shared across all 25 comparisons) (Figure 2A). A comparison of the TEs that were annotated as structurally intact or homology-based revealed that structurally annotated TEs are depleted for core and near-core features, exhibiting a higher frequency of variable and private elements (Figure 2B). Non-syntenic genes exhibit a distribution of classifications that are quite similar to the homology-annotated TEs, while the syntenic genes are enriched for core features with relatively few variable and private annotated features (Figure 2B). This is consistent with the observation that, in maize, many non-syntenic genes exist within or substantially overlap TEs (26).

**Figure 2.**
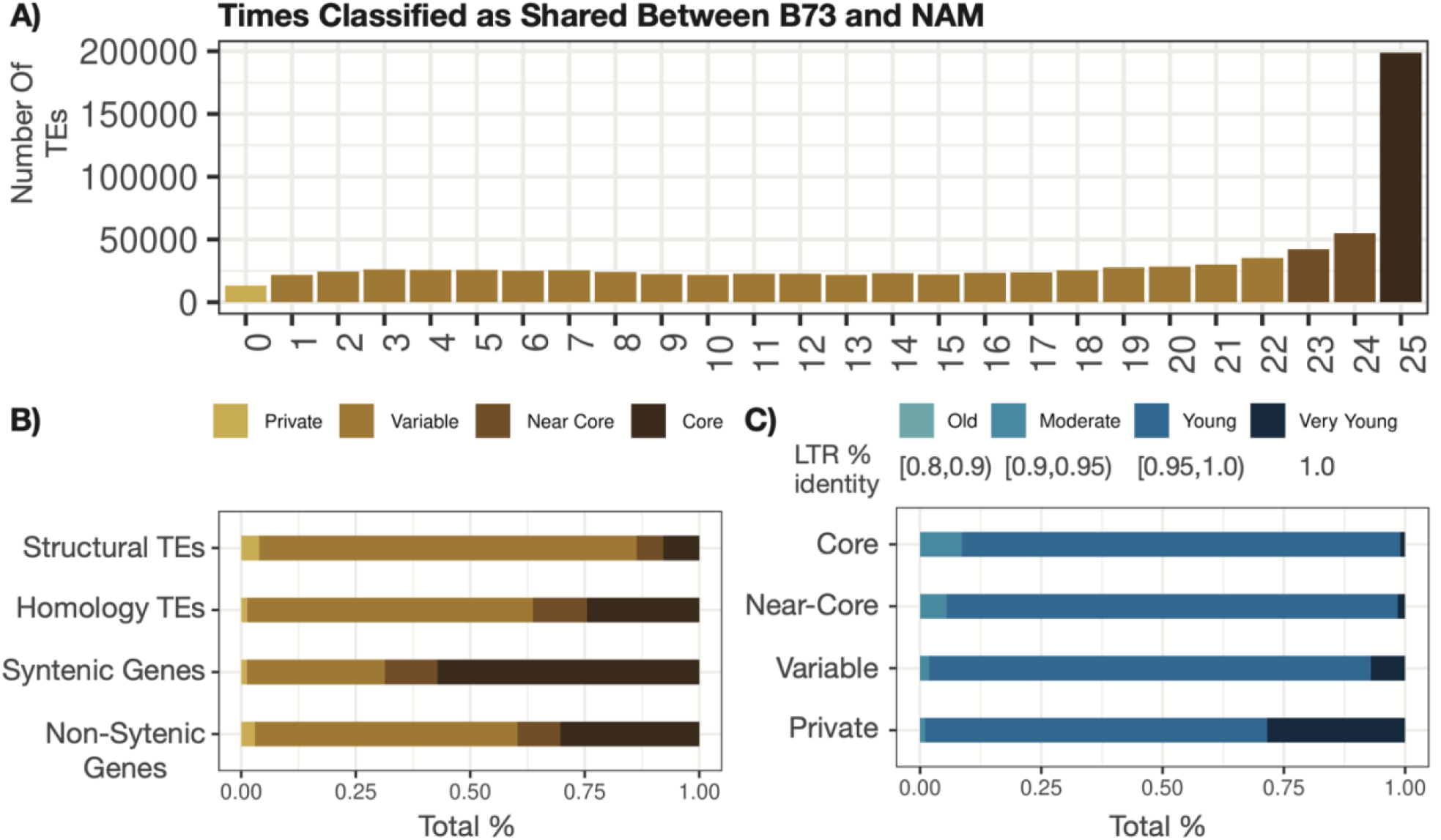
Consistency of B73 Feature Classifications Across B73 versus NAM comparisons. Every feature in the genome (i.e., TE or gene annotation) is classified as shared, polymorphic, or ambiguous and binned into either core (classified as shared across all 25 comparisons – dark brown), near-core (classified as shared in 23 – 24 comparisons – brown), variable (classified as shared in 1 – 24 comparisons – light brown), or private (never classified as shared across all comparisons – tan). (A) Distribution of all B73 TEs across all NAM comparisons. (B) The proportion of B73 structurally-annotated TEs, homology-annotated TEs, syntenic genes, and non-syntenic genes classified as either core, near-core, variable, or private. (C) The age of a structurally-annotated LTR was estimated from the percent identity between the two long terminal repeats and binned into either very young (% identity = 1), young (% identity within [0.95,1)), moderate (% identity within [0.9,0.95)), and old [0.8,0.9)) with darker colors representing younger TEs. B73 structural TEs were partitioned depending on whether they are core, near-core, variable, or private and the proportion of each category classified as young, very young, moderate, or old is shown.

Structurally annotated LTR elements could be used to assess the relative age of TEs that were classified as either core or variable. The two long terminal repeats are 100% identical at the time of insertion due to the mechanism of LTR TE mobilization. Older elements will accumulate polymorphisms that can distinguish the two long terminal repeats providing a proxy for increasing age of each element. Structurally annotated LTR elements that are private to B73 relative to the other NAM genomes have substantially more very young (LTRs are 100% identical) relative to core and near-core LTR elements suggesting that at least a subset of private elements may represent relatively recent insertions (Figure 2C).

### Different types of TEs exhibit variable polymorphic frequencies

We proceeded to investigate whether there are differences in the frequency of polymorphic TEs for different superfamilies of transposable elements. TEs can be subdivided based on their annotation method (structural vs homology) as well as their superfamily. There are four major superfamilies of class I elements — LINE (RIL), LTR-Copia (RLC), LTR-Ty3 (RLG), and LTR-Unknown (RLX) — and five superfamilies of class II elements — DTA, DTC, DTH, DTM, and DTT based on homology to the hAT, CACTA, Pif/Harbinger, Mutator, or Tc1/Mariner superfamilies respectively. The retrotransposon superfamilies (RLC, RLG, and RLX) do not show substantial variation in the proportion of elements that are classified as polymorphic and have similar frequencies for both homology and structural annotations. In contrast, class II TIR transposon superfamilies exhibit more variable frequencies of polymorphic TEs (Figure 3A). Both homology and structural annotations of DTT elements exhibit relatively low (<20%) frequencies of polymorphic calls. The structural annotations of the other superfamilies are much more likely to be polymorphic with nearly 50% of structural DTA elements classified as polymorphic. It is unclear whether these differences are due to biological differences among superfamilies or technical artifacts such as differences in sizes or annotations.

**Figure 3.**
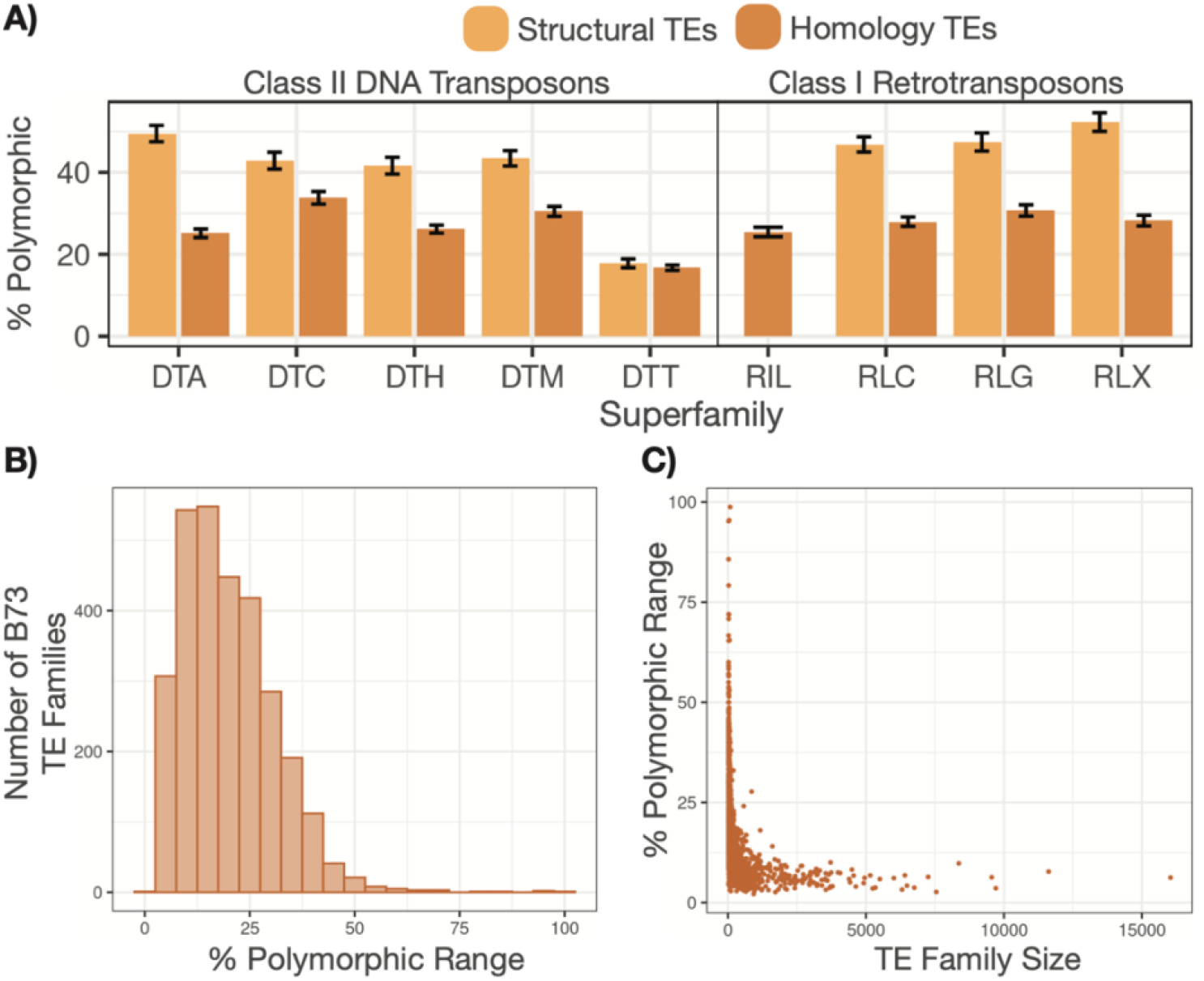
Distribution of polymorphic B73 TEs across superfamilies and within distinct families. (A) B73 TEs were grouped both by their assigned superfamily (x-axis) and by whether they were a structurally-annotated (yellow) or homology-annotated (orange) element. Within a specific B73 versus NAM comparison, the percent of those elements classified as polymorphic was calculated. The height of the bar shows the mean percent across all 25 comparisons with error bars showing the standard deviation. (B) For each B73 TE family with greater than 20 copies in the genome, we determined what percent of that family was classified as polymorphic in a given comparison. We then plotted a histogram showing the range of these percent polymorphic values across all 25 comparisons. (C) The x-axis shows the number of copies for B73 TE family (> 20) in the B73 genome, while the y-axis shows the percept polymorphic range for that family.

TE superfamilies can be further subdivided into families, which contain related elements. In order to assess whether there are specific TE families with high levels of polymorphic elements, we also did a per-family analysis of the frequency of polymorphic elements. Each TE family was assessed in 25 total contrasts between B73 and each NAM genome providing multiple estimates of the frequency of polymorphic elements within a family. The range of the percent of polymorphic elements for a given family provides insight into whether that family has particularly high or low frequencies of polymorphic elements in one or a handful of contrasts. We limit this analysis to families with at least 20 members in B73. We find substantial variability in the proportion of polymorphic elements for small families (Figure 3B). However, the larger families tend to have a constrained proportion of polymorphic TEs (Figure 3C). In addition, there is limited evidence for outlier genomes in these contrasts (Figure 3B-C). This suggests that there are few examples of specific TE families that have increased substantially in copy number in B73 relative to any of the other genotypes or vice versa.

#### Polymorphic TEs are often located within larger structural variants

There are many thousands of polymorphic TEs identified in any genome comparison in maize. It can be tempting to think of these as representing TE insertion polymorphisms. However, visualization of specific chromosomal regions revealed that in many cases there are structural variants (SVs) that include multiple TEs from distinct superfamilies as well as non-TE sequence (Figure 4A-B). A careful examination of one of these SVs that is present in 14 of the NAM genomes reveals two TE fragments that are polymorphic and two TEs at the edges that are ambiguous due to the boundaries of this structural variant falling within annotated TEs (Figure 4A-B). These likely represent deletion events that removed TEs present in the ancestral sequence. Instead of solely focusing on a TE-centric analysis of variation as described above, we characterized the full set of large (> 50bp) structural variants to understand how these SVs may be associated with transposon sequence(s) (Figure 4C).

**Figure 4.**
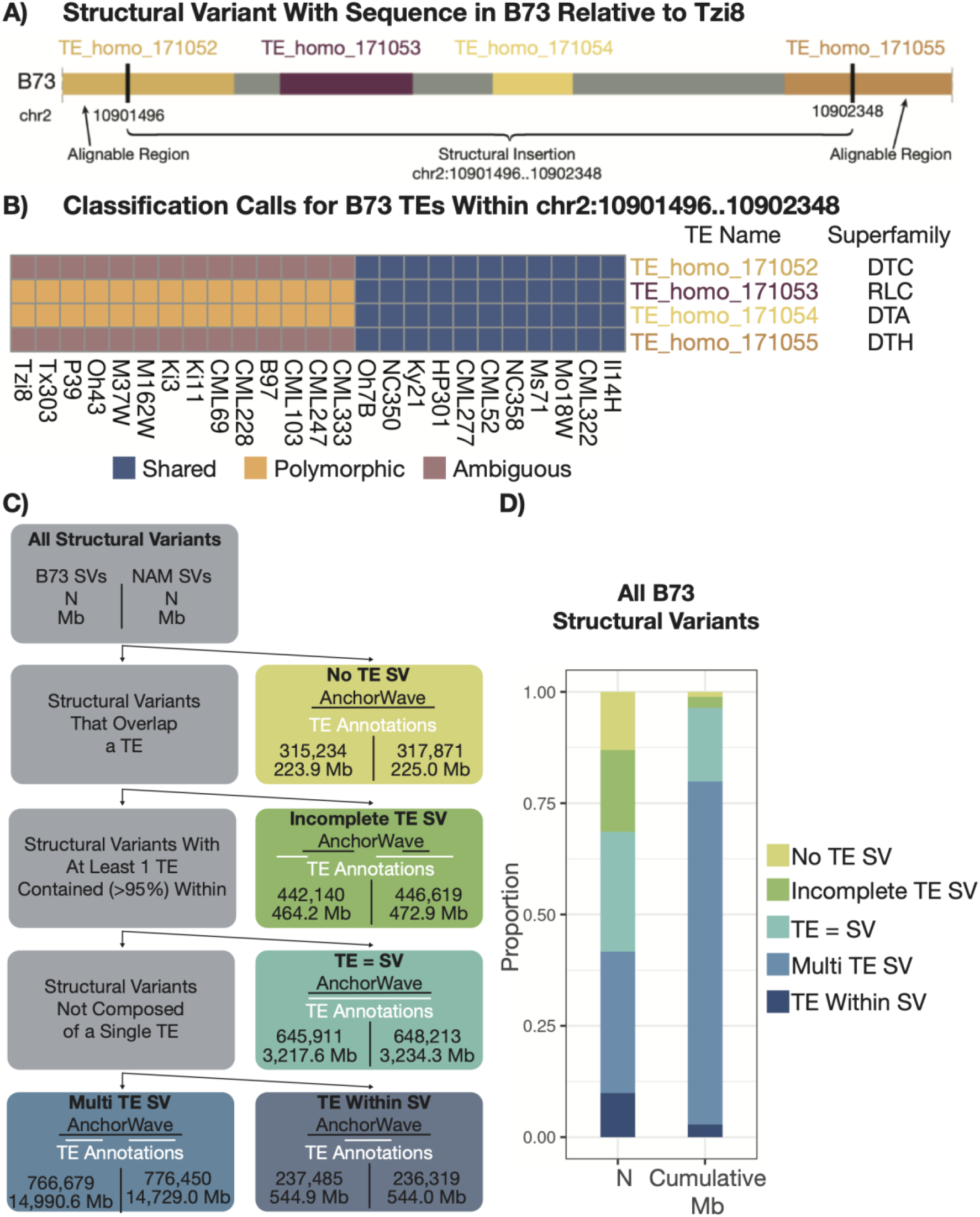
Structural Variants Exhibit Differential TE Content. (A) A visualization of a SV sequence in B73 relative to Tzi8 that is flanked by alignable regions. TE annotations in this region are overlaid with different colors representing the superfamily of the TE. (B) A heatmap showing the classification calls for the B73 TEs present in the SV shown in (A) across all of the NAM genomes. Colors represent whether the TE was classified as shared (blue), polymorphic (yellow), or ambiguous (purple) in that comparison. Columns are clustered based on similarity of alignable proportions of visualized TEs and show two distinct haplotypes where the SV is present (left) or absent (right). (C) A flowchart showing how SVs were classified depending on their overlapping TE content. (D) A stacked barchart showing the proportion of SVs (in terms of total number – left column – and cumulative Mb – right column) with sequence in B73 relative to another NAM genome with colors delineating the different classes of SVs characterized in (C).

In the analysis of all NAM genomes compared to B73, there were a total of > 2.4 million SVs (> 50bp) that represented sequence present in B73 absent in a NAM comparison that cumulatively accounts for 19,441 Mb of sequence (Figure 4C-D). These exhibit a wide spectrum of variation for the size of the SV sequence with 54.7% that were > 1kb, 22.7% that were greater than >10kb, and 0.43% that were greater than >100kb. An analysis of the TE annotations within the SV sequences revealed that 13% of the SV sequences in B73 do not overlap any TE sequence, and these SVs are classified as ‘No TE SV’. These ‘No TE SV’s tend to be relatively small and only account for 1.2% of the total Mb of SV sequence in B73. There were another 18.4% of SV sequences (2.4% of SV sequence Mbs) that only have partially (<95%) overlapping TE sequences, which were classified as ‘Incomplete TE SV’. These include examples in which only one edge of the TE overlaps the SV sequence as well as instances of SV sequences that are located entirely within a longer TE sequence. These ‘Incomplete TE SV’ overlaps likely represent deletion events that removed either one edge of the TE or an internal portion of the TE in the non-B73 genome.

The remaining 68% of SV sequences include at least one TE that was > 95% contained within the SV. These were divided into three further groups. The ‘TE = SV’ group is limited to instances in which both > 95% of the TE overlaps the SV and > 95% of the SV overlaps the TE. These account for 26.8% of SVs and 16.6% of the total Mbs of SVs. These ‘TE = SV’s likely represent TE insertion polymorphisms as well as some potential TE excision events. The other two groups that contain at least one full (> 95%)TE were divided based on whether they included multiple full length TEs (‘Multi TE SVs’) or only a single full TE that accounted for less than 95% of the SV sequence (‘TE Within SV’). The ‘Multi TE SV’s make up 31.8% of SVs and account for 77.1% of the Mb of SV sequence, of which 84% is actually TE sequence. The ‘TE within SV’ category accounts for the remaining 9.8% of SVs and 2.8% of the Mbs of SV sequence. The same analyses were also performed for the SVs that are present in the NAM genomes and absent in B73 and reveal similar trends as B73 present sequences (Figure 4C).

The classification of SVs relative to TE annotations revealed that, on average, 85% of the total SV sequence in B73 compared to any one NAM genome is annotated as TE. This supports the hypothesis that TEs are a major contributor to structural variation among maize genomes. However, while TE sequence is a predominant component of SVs, it seems that canonical insertion polymorphisms (represented by the ‘TE = SV’ category) only account for a fraction of the SVs. Our TE centric comparison of genomes identified a total of approximately 6.47 million B73 polymorphic TE classifications across all comparisons of B73 to the other NAM genomes with a mean of 258,926 B73 polymorphic TEs in a given comparison to any one of the NAM genomes. The vast majority (99.9%) of these polymorphic TEs have >95% overlap with a single SV. These polymorphic TEs could be assigned to the ‘TE = SV’ (671,219 – 10.4%), ‘Multi TE SV’ (5,556,116 – 85.9%), and ‘TE Within SV’ (237,432 – 3.7%) groups. Overall, these analyses suggest that the majority of polymorphic TEs are not necessarily due to true TE insertion polymorphisms but are instead due in large part to deletion events that remove TE sequence.

### Polymorphic TEs within SNP depleted blocks

Genomic regions that are highly alignable with very low SNP rates when comparing B73 to another NAM genome likely reflect chromosomal regions recently derived from a common ancestor. In fact, relatively long (> 1Mb) SNP depleted regions are likely diverged for only tens to hundreds of generations. Using the pairwise AnchorWave alignments, we sought to identify these large SNP depleted regions in order to use them to monitor relatively recent changes in TE content in the NAM genomes (Figure S3). Our analysis was restricted to regions of at least 2Mb with a SNP rate that was at least 100-fold lower than the genome-wide average SNP rate (see methods for details). There were a total of 213 of these SNP depleted regions that were identified based on comparisons of B73 to all of the other 25 NAM inbred parents (Table S2). There were no SNP depleted regions identified in comparisons of B73 and four of the NAM genomes (CML69, CML247, CML277, and NC350). In contrast, > 20 regions were identified between B73 and each of B97, Ky21, MS71, Oh7B, and Oh43. The size of the regions was quite variable (Figure S4). While many (26) of the regions were only 2Mb in length in B73, there are also 23 SNP depleted regions that are at least 10Mb in length. These 213 SNP depleted regions account for approximately 1,048Mb cumulative base pairs of the B73 genome.

We expect to observe little variation in the SNP depleted regions given the relatively short divergence time between the two pairs of haplotypes. The percent of Mb attributable to structural variant sequences in SNP depleted regions is greatly reduced in comparison to the genome wide percentage (Figure 5A), and we find highly similar TE content as well. The B73 haplotypes in these regions contain 424,665 TEs (including 31,230 structurally annotated TEs). The vast majority (> 99.9%) of these TEs were classified as shared (Figure 5B). There were 410 B73 TEs in these regions classified as polymorphic and another 110 TEs classified as ambiguous. A similar analysis of the 424,577 TEs annotated in the NAM haplotype sequences for these regions revealed 576 polymorphic and 127 ambiguous TEs. The 986 polymorphic Tes within the SNP depleted regions likely represent a combination of novel insertions as well as deletion and/or excision events that could remove TE sequences. The analysis of SVs within these regions identified a total of 690 SVs including 139 ‘TE = SV’ putative insertions in B73 and 79 ‘TE = SV’ putative insertions in the non-B73 genomes (Figure 5C-D).

**Figure 5.**
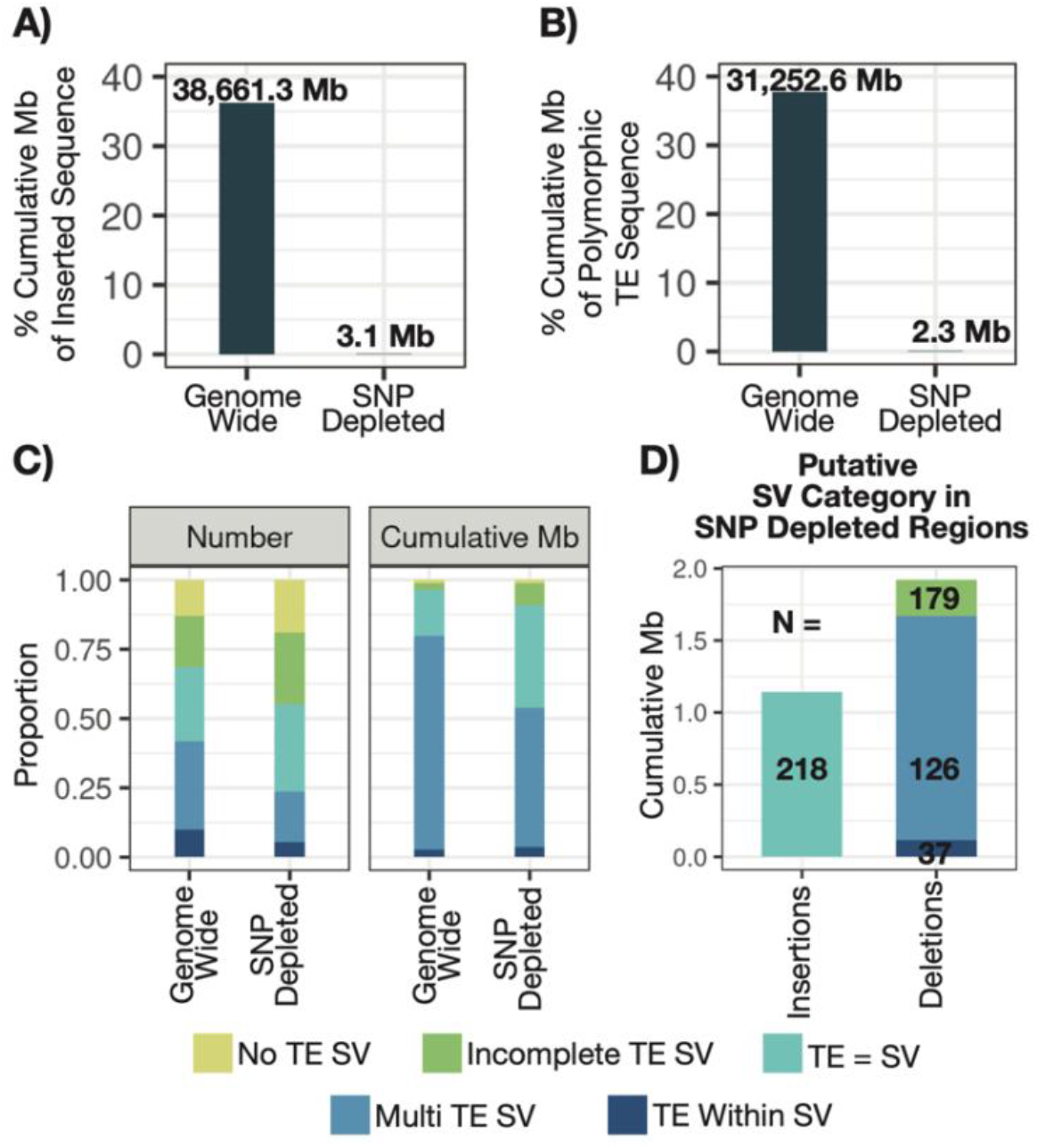
Structural and TE Variation in SNP Depleted Regions. (A) The percent cumulative Mb of inserted sequence from structural variants combined across B73 and NAM comparisons both genome-wide (left) and in SNP depleted regions (right) out of the total cumulative Mb of genomic sequence. (B) The percent cumulative Mb of polymorphic TE sequence present in B73 and NAM comparisons both genome-wide (left) and in SNP depleted regions (right) out of the total cumulative Mb of TE sequence. (C) A stacked barchart showing the proportion of structural variants (in terms of total number – left panel – and cumulative Mb – right panel) with sequence in one genotype relative to another across our B73 to NAM comparisons with colors delineating the different classes of SVs. The proportion is shown both genome wide (left) and in SNP depleted regions (right). (D) SV groupings can be further reduced into either putative insertions (left), comprised of ‘TE = SV’ events, or putative deletions (right), comprised of ‘Incomplete TE SV’, ‘Multi TE SV’, and ‘TE Within SV’ events. A barchart shows the cumulative Mb of these putative SV categories in SNP depleted regions.

Only 224 of the 986 polymorphic TEs were classified as being located within ‘TE = SV’ variants and likely represent insertion events. The majority of the remaining 762 polymorphic TEs are located within ‘Multi TE SV’ (725 TEs) or ‘TE Within SV’ (37 TEs) variants and likely represent recent deletions that have removed TEs. The 224 ‘TE = SV’ polymorphic TEs were candidates for relatively recent insertion polymorphisms from active TE families. These 224 TEs were from many different superfamilies of TEs and had relatively balanced numbers of insertions in the NAM genomes with no evidence for massive burst in any particular genome (Figure S5, Table S3). A per-family analysis revealed four TE families that account for > 10 ‘TE = SV’ polymorphisms within the SNP depleted regions, and, in total, these four families accounted for nearly 60% of all ‘TE = SV’ events in these regions (Figure 6A). These include 67 members of the family CRM2_7577nt, 34 members of the family DTA_ZM00383, 17 members of the family ji_AC204382, and 15 members of the family ji_AC215728. We characterized several features about these families with > 10 polymorphic ‘TE = SV’ events. In each case, there are roughly similar numbers of insertions in B73 and each of the other NAM genomes, and we find evidence for insertions in multiple genomic regions (Figure S6).

**Figure 6.**
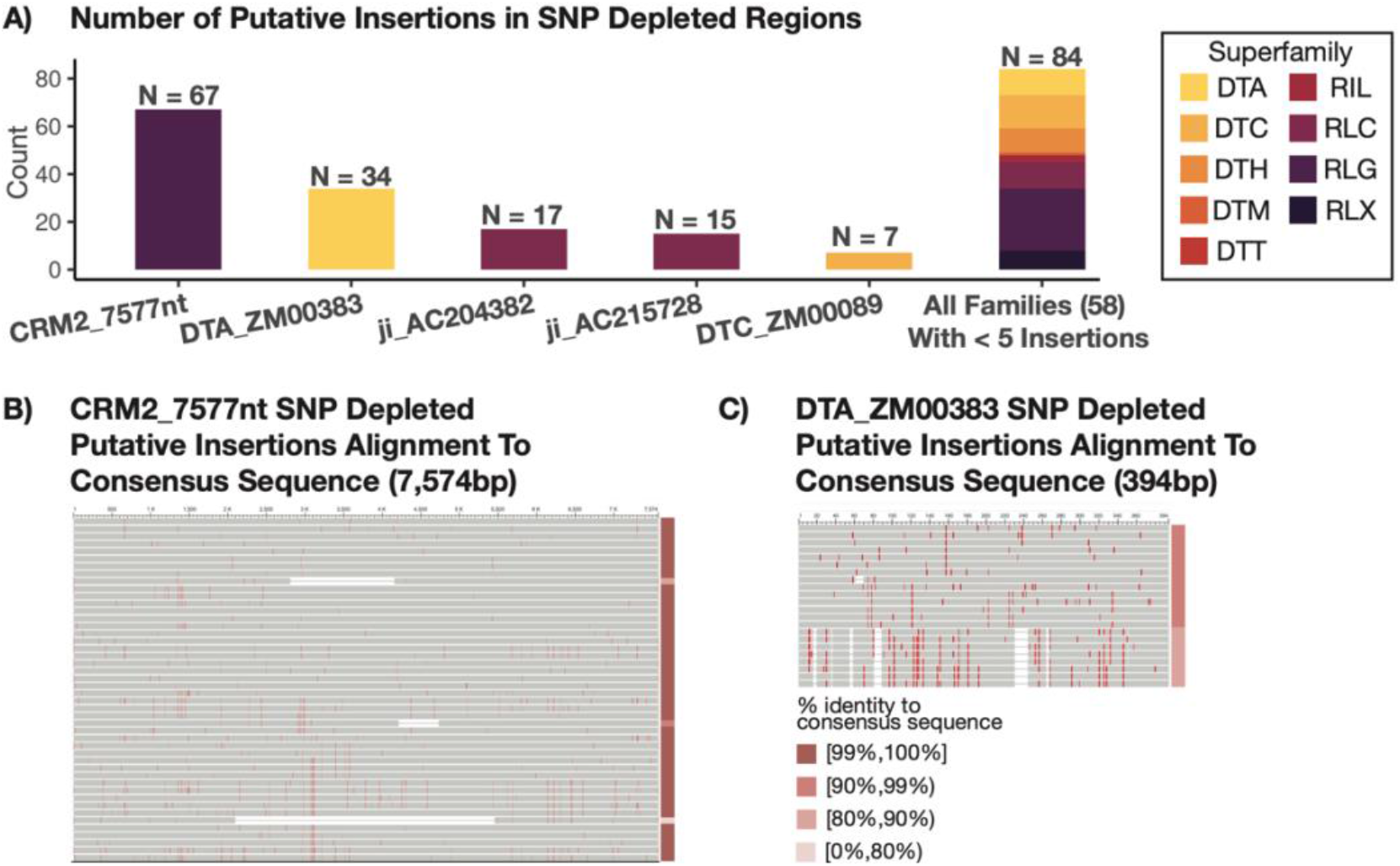
Putative Insertions in SNP Depleted Regions Highlight Potentially Active TE Families. (A) Putative insertions in SNP depleted regions were grouped by family to assess whether specific families show higher insertion events, suggesting potential transposition activity. A barchart shows the number of putative insertions for the five TE families with greater than five insertion events in SNP depleted regions, as well as a stacked bar showing the cumulative count of the remaining 58 families all of which have less than five insertion events. The specific number of events is annotated above each bar. Color indicates the superfamily classification for that TE family. Putative insertions (excluding solo elements) in SNP depleted regions belonging to the TE family CRM2_7577nt (B) and DTA_ZM00383 (C) were aligned to the consensus sequence for that family used in the annotation process. Vertical red lines indicate the presence of SNP relative to the consensus sequence. The percent identity to the consensus sequence is represented using color in the bar to the right of the multisequence alignment with darker pink colors representing a greater percent identity to the consensus sequence.

Each of these families exhibit evidence that a specific subset of elements might be active. For example, the majority of the CRM2_7577nt insertions (61 of 67) are between 7500bp and 7600bp with ten insertions each that are exactly 7577 or 7581 base pairs in length (Figure S6). Similarly, all putative insertions in SNP depleted regions belonging to the family DTA_ZM00383 are between 395bp and 406bp with 21 of 34 members being exactly 395bp in length (Figure S6). At a per-family level, sequence similarity between the putative insertions in SNP depleted regions to the consensus sequence used in the annotation process was determined (Figure 6B-C,S7). These putative insertions show high sequence similarity across all four families. With the exceptions of TE sequences with small InDels present, putative insertions in SNP depleted regions for CRM2_7577nt all show greater than 99% identity to the consensus sequence (Figure 6B). Putative insertions for DTA_ZM00383 in SNP depleted regions similarly show high sequence similarity to the consensus, and SNP content amongst these insertions suggest two to three elements may have been actively proliferating (Figure 6C). Genome-wide analysis of the members of these families revealed substantially more variation in the size of all elements in these families which suggests that only a subset of these elements might be capable of movement.

While the analysis of the SNP-depleted regions provides an opportunity to capture evidence for recent transposition events, it is likely that recent events are occurring genome-wide. The analysis of all ‘TE = SV’ polymorphic TEs revealed that these four families each have high levels of ‘TE = SV’ events. There are 8,307, 6,945, 11,481, and 17,599 events respectively for the families of CRM2_7577nt, DTA_ZM00383, ji_AC204382, and ji_AC215728. Among the 779 TE families with an average of > 100 copies per genome, these four families are all in the top 25% for proportion of ‘TE = SV’ events.

## Discussion

As researchers have moved from the era of single reference genomes for any species to producing multiple high-copy genome assemblies for genetically diverse individuals of a species, substantial variation in genome content has been observed (23,34). We were particularly interested in studying the contribution of transposable elements (TEs) to this genomic variation. We sought to understand both the contribution of TEs to structural variants and to understand the relative dynamics of TE insertions compared to deletion events that might contract genome size via removal of TEs and other sequences.

The high-quality NAM genomes provide an opportunity to study these dynamics (23). The TEs within these genomes were all annotated using a consistent approach (31). From these annotations, we were able to classify the shared or polymorphic status for the majority of the elements in the genome. These classifications relied upon the precise whole-genome alignments generated by AnchorWave (30). We found that there are abundant polymorphic TEs in the comparison of maize genomes. There are approximately 900,000 annotated TEs in any NAM genome. Across our comparisons, we found that ∼650,000 of these TEs are shared between B73 and any NAM genome while the remaining ∼250,000 TEs are polymorphic. These results are quite similar to prior work that assessed shared and polymorphic TE content in four genomes using a different approach for annotation and classification (3,26,35). This leads to substantial variation among the genomes in terms of the presence of TEs at specific genomic regions and results in many genes being located near polymorphic TEs. Only a subset of maize TEs are shared among all the NAM genomes leading to highly variable haplotype structures.

While it can be quite useful to classify TEs as shared or polymorphic, the actual haplotype structures reveal more complex patterns. Often polymorphic TEs are part of a larger structural variant that includes multiple TEs. This can complicate functional analyses as it becomes impossible to monitor the potential effect of just one polymorphic TE as there are multiple TEs all in complete linkage. We therefore implemented a strategy to classify all structural variants (InDels > 50bp) in relation to TE annotations. This revealed that many (85.8%) of the polymorphic TEs in B73 actually occur as part of ‘Multi TE SV’ events. Only a subset (10.4%) of these polymorphic TEs are instances in which the boundaries of the TE and the SV are quite similar (‘TE = SV’ events).

We infer that the majority of the ‘Incomplete TE SV’, ‘Multi TE SV’, and ‘TE Within SV’ events are likely deletions influencing ancestral TEs rather than polymorphic insertions. This inference is further supported by the fact that many of these SVs have edges that occur within the middle of an annotated TE in the genome that lacks the deletion. It is certainly true that some of the events may be insertion polymorphisms with nested TEs or other structures, but the majority are likely deletion events. The ‘TE = SV’ events are more likely to represent insertional polymorphisms, although some could represent TE excisions, especially for terminal inverted repeat DNA transposons.

Maize is an outcrossing species and several of the NAM inbreds have pedigrees that include common parents. This leads to the presence of large SNP depleted regions that likely represent inheritance of an extended haplotype from a common parent in the past 10s to 100s of generations. These regions provide an opportunity to study the ongoing dynamics of structural variation and TE polymorphisms in modern maize inbreds. While these regions have very few SNPs as expected for common descent, there are still some examples of structural variants and TE polymorphisms. It is worth noting that >99.9% of the annotated TEs within these regions are classified as shared suggesting that the ongoing movement of TEs is quite limited. However, the rare polymorphisms within these regions provide insights int0 the current expansion and contraction of TEs. Only 224 (23%) of the 986 polymorphic TEs in SNP depleted regions represent ‘TE = SV’ events. The majority of the polymorphic TEs likely occur as a result of deletion events rather than novel insertions. In these regions, there are a total of X Mb of putative deletions (‘Incomplete TE SV’, ‘Multi TE SV’, and ‘TE Within SV’) and X Mb of putative TE insertions (‘TE = SV’ events). This suggests that the maize genome may be in a phase of contraction rather than expansion as TE removal is currently outpacing TE insertions.

The characterization of the ‘TE = SV’ events within these SNP depleted regions identified several TE families with potential transposition activity in modern germplasm. These included several LTR families (CRM2_7577nt, ji_AC204382, and ji_AC215728) and a DNA TIR family (DTA_ZM00383). We did not find evidence that these families were only active in particular genotypes but instead see likely evidence for low rates of movement across multiple genotypes. There do seem to be particular members of these families that are mobile as many of the putative insertions are of quite similar length and sequence identity. Genome-wide analyses suggest high levels of polymorphic TEs within these families. Further studies to characterize these potentially active TEs will be important in characterizing potential autonomous elements and the impacts of these TEs.

Combined our results highlight how both insertions and deletions of transposable elements contribute to structural variation in maize. It is compelling to think of TEs as generating insertions. Both their mechanisms of movement and characterization as ‘selfish’ genetic elements evoke the idea of sequence gain. However, unmanaged TE proliferation would lead to genomic bloat as genomes become riddled with TEs. While genomes have evolved myriad mechanisms to silence TE proliferation, ongoing deletion events provide another strategy for mitigating the spread of TEs. Genome-wide deletion rates are likely governed by factors independent of TE transposition rates, but TE proliferation could facilitate ectopic recombination events between similar TE sequences, resulting in a deletion event. Further investigation into the insertion and deletion rates of TEs is warranted to better understand how TEs influence genomic variation.

## Methods

### Characterization of structural variation among NAM genomes

High-quality genomic sequences have been produced for the 26 NAM inbred founder lines (23). AnchorWave v1.0.1, a recently developed approach used to perform pairwise whole-genome alignments, was used to compare each of the NAM inbred genomes to B73 (30) via the ‘genoAli’ command and ‘-IV’ parameter. The MAFToGVCF plugin of tassel v5.2.82 (36) was used to reformat genome alignments in MAF format into variant calling records in GVCF format. The resulting GVCF files contain records of all nonvariant and variant sites, including single nucleotide and structural differences. Given our interest in shared or polymorphic TEs, we condensed the GVCF output by combing the nonvariant sites, single nucleotide variants, and small (<50bp) insertion/deletion variants into a single class of ‘alignable’ regions. ‘Structural variants’ in one genotype relative to another were defined as regions > 50bp for which the size in the other genotype is 0bp. The remaining variants all include at least one base pair in each genotype that is not fully aligned, and these regions were consequently classified as ‘unalignable’. Gaps in the AnchorWave alignment were identified and classified as ‘missing data’, but these regions were treated as unalignable in all downstream analyses.

### TE annotation processing

The TE content for each of the NAM genomes had been previously annotated using panEDTA (23,31,32)[14,25,26]. This process initially identifies TEs based on structural features using tools including LTR_Finder (37), LTRharvest (38), and TIR-Learner (39). These structurally identified TEs are then used to create a panTE library across all of the genomes that is used to perform homology based annotation of non-structurally intact TE fragments. We filtered out annotations for non-TE repeats, helitrons, and specific features of structurally annotated LTRs (e.g., target site duplicataions, long terminal repeats) such that we only retain the full-length structural annotation. We removed a handful of ‘duplicate’ annotations with different IDs but identical coordinate positions, superfamily classifications, and family classifications. With duplicate annotations, we prioritized retaining structural annotations over homology annotations. If both annotations were identified using the same method, we randomly decided which one to keep.

Preliminary analyses of the previously released output using bedtools (v2.30.0) (40) identified potentially problematic overlapping annotations. One family (‘DTA_ZM00081_consensus’) frequently had homology annotations that occurred in multiple regions throughout LTR elements suggesting potential contamination of this TIR element with LTR related sequences in the original Maize TE Consortium (MTEC) library that carried forward into the panEDTA library. Therefore, all annotations with this family ID were removed from downstream analyses. We also identified examples of overlapping TE annotations that seemed biologically unfeasible. TE annotations whose start or end position was within 5 base pairs of the start position or within 5 base pairs of the end position of a structurally annotated TE were filtered out. If two structural annotations overlapped in this way, we prioiritized retention of the larger structural annotation and randomly selected one for retention if they were equally sized. Any homology TE annotation that had greater than 10% overlap (including those that were contained with 100% overlap) with another homology annotation was removed. Additionally, any homology TE annotation that overlapped a structural TE annotation by greater than 5% but was not fully contained within that structural annotation was removed. Finally, we filtered out structural TE annotations that overlapped but were not contained within another structural TE annotation. We prioritized retention of Class II annotations and longer annotated elements when deciding which structural annotation to keep. This resulted in a final annotation in which tehre are very few examples of the same region being annotated as part of multiple TE features and allows for a more accurate assessment of the TE bae pairs within each of the genomes.

### Gene annotation processing

Annotated genes for each NAM line were characterized and similarly obtained from Hufford et al. (2021). As part of this work, each gene was classified as either syntenic or non-syntenic relative to sorghum. Annotations could be further broken down into exon-only or full-length annotations.

### Identification of shared and polymorphic genomic features

The processed AnchorWave GVCF files allowed classification of all segments of pairwise contrasted genomes into alignable, inserted, or unalignable sequence. The genome-wide annotations for transposable elements or genes were then intersected with these regions using bedtools (v2.30.0). Any features that had at least 95% of its sequence overlapping alignable regions was classified as a shared TE between the contrasted genomes. Features were classified as polymorphic if they had at least 95% sequence overlap with a structural variant in their own lineage (i.e., B73 TEs were polymorphic if they had 95% overlap with structural variants with sequence in B73). The remaining features that did not include at least 95% overlap with either alignable or structural variant sequence were classified as ambiguous, as their exact status (i.e., shared or polymorphic) could not be confidently determined.

We classified both the filtered EDTA TE annotations and the canonical gene annotations across the NAM genomes as either shared, polymorphic, or ambiguous. Canonical gene annotations were classified using either exon-only coordinates or full-length coordinates for each model. Due to the pairwise nature of the AnchorWave alignments, the B73 TE annotations were classified 25 times, one for each query comparison, while the TE annotations for the remaining 25 NAM genomes were only classified once in relation to their presence or absence in B73.

### Identification of SNP depleted regions

Between B73 and each of the NAM genomes, the number of SNPs and amount of alignable sequence called using the AnchorWave alignment was used to identify SNP depleted regions in each pairwise comparison. We used a sliding window approach to count the number of SNPs and base pairs of alignable sequence in 1Mb windows offset by 250kb from the start to end of each chromosome. Normalized SNP counts for each 1Mb window were then determined by dividing the SNP count by the total amount of alignable sequence in the 1Mb window. We identified the subset of 1Mb windows that had >950,000 base pairs of alignable sequence and had a normalized SNP rate less than 1 in 10,000 (the average SNP rate was 1 in 44). We further required a minimum of 5 consecutive 1Mb sliding windows that meet these criteria in order to identify at least 2Mb regions in pairwise comparisons that were highly depleted of SNPs and likely represent identity by state. For all analyses, we offset the start and end coordinates for each SNP depleted region by 100,000 base pairs to ensure the boundaries of the region did not extend beyond the putative identity by state region of the genome.

### Evaluation relatedness of putative insertions in SNP depleted regions

Four TE families had several putative insertions within SNP depleted regions – CRM2_7577nt, DTA_ZM00383, ji_AC2043821, and ji_AC215728. Within each family of interest, all TE sequences identified as putative insertions in SNP depleted regions across all of the pairwise comparisons were extracted. Any solo elements were dropped from further analysis. To determine the phylogenetic relationship between copies within each family, the remaining sequences were aligned with MUSCLE using default settings (41). These aligned sequences were then trimmed using trimAL with paramters ‘-automated1’ (42). Trimmed sequences were then aligned again using MUSCLE and default parameters. A phylogenetic tree was generated for eacfh family using RAxML with settings ‘-f a -m GTRGAMMA -p 12345 -x 12345 -# autoMRE’ (43). Phylogenetic trees were plotted with the ggtree R package (44).

## Declarations

## Ethics approval and consent to participate

Not applicable

## Consent for publication

Not applicable

## Availability of data and materials

All datasets used in this analysis are publicly available. TE annotations, gene annotations, and gene synteny calls were downloaded from MaizeGDB. TE and gene annotations were downloaded from https://maizegdb.org/NAM_project, while synteny classifications for the NAM genes were download from https://ars-usda.app.box.com/v/maizegdb-public/folder/186350887665. All remaining datasets have been deposited on Dryad and can be accessed at https://doi.org/doi:10.5061/dryad.5qfttdz9t. This includes the pairwise AnchorWave alignments in both MAF and GVCF format, TE and gene classification calls, and the sliding window SNP counts used to identify SNP depleted regions. Scripts used to filter and analyze data are available on GitHub at https://github.com/mam737/PolymorphicTEs_NAM. To visualize pairwise alignments with overlapping TE and gene annotations, an R Shiny App Web Browser was developed and can be found at https://mmunasin.shinyapps.io/nam_sv/

## Author Contributions

M.M. and N.S. conceived the study and designed the logical framework used throughout. M.S. and B.S. generated the pairwise AnchorWave alignments. M.M. wrote all scripts used to filter and summarize relevant datasets, classify features and structural variants, and visualize results with assistance and feedback from all authors. A.R., C.M., K.M., C.H., and Y.B. provided assistance in accessing and analyzing relevant datasets. C.M. ran analyses evaluating the relatedness of putative insertions in SNP depleted regions. M.M. and N.S. wrote the first draft and all authors contributed to the writing of the manuscript. N.S. and Y.B. supervised the project.

## Acknowledgements

We would like to thank Shujun Ou for sharing the panEDTA TE annotations with us directly. We would also like to thank Jeff Ross-Ibarra, Emily Josephs, Nathan Catlin, Zach Myers, Erika Magnusson, Cathy Rushworth, and Shelley Sianta for useful comments, insights, and suggestions throughout the analysis. Finally, we thank the Minnesota Supercomputing Institute at the University of Minnesota (https://www.msi.umn.edu) for providing resources that contributed to the research results reported within this article.

