## Supplemental Figures for "Combined analysis of transposable elements and structural variation in maize genomes reveals genome contraction outpaces expansion"

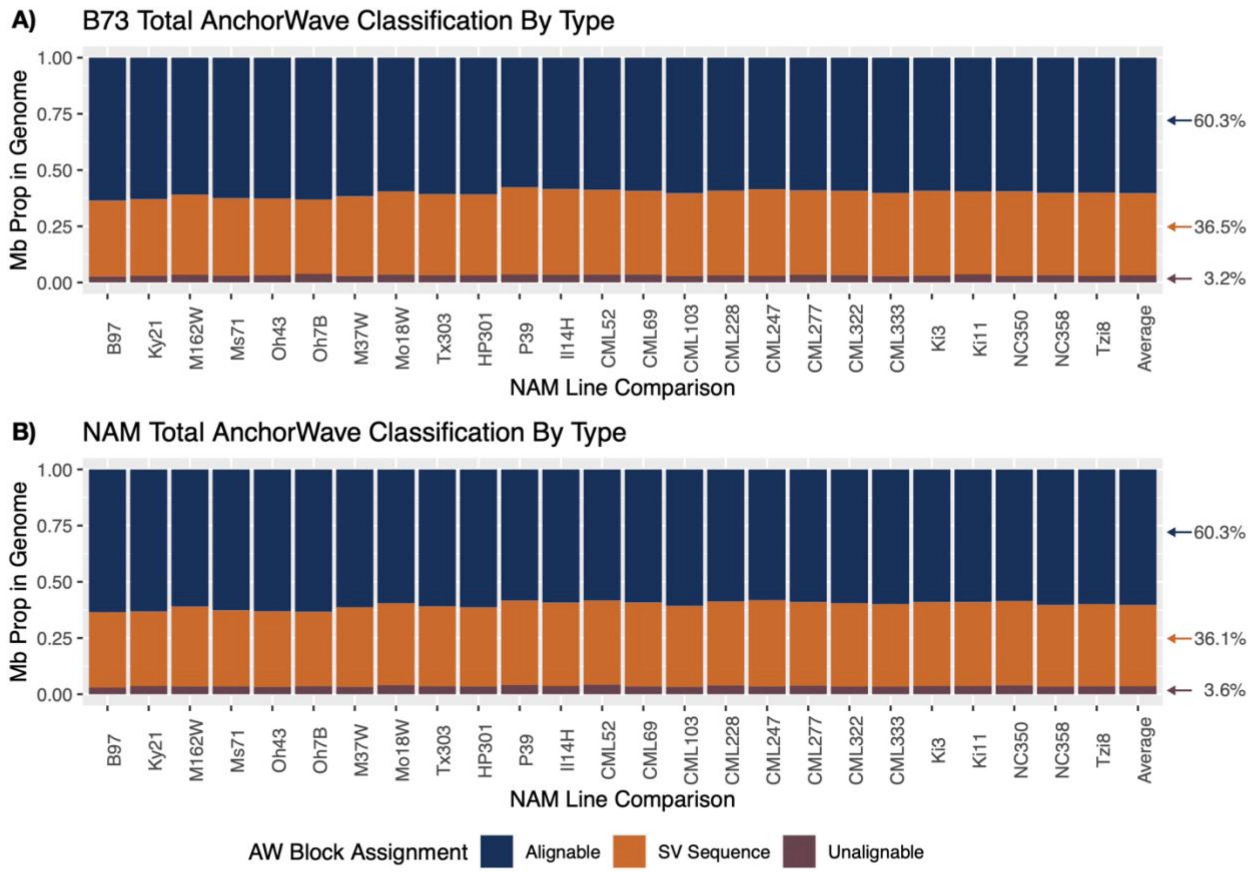


**Figure S1 Proportions of AnchorWave Classifications Across NAM Lines.** For each pairwise contrast between B73 and a NAM genome, each region of the pairwise AnchorWave alignment could be classified as either alignable (dark blue), structural variant sequence (orange), or unalignable (purple). (A) Barplots showing the proportion in Mb of the B73 genome classified as each group against every NAM line (x-axis). (B) Barplots showing the proportion in Mb of every NAM genome (x-axis) classified as each group against B73. The last bar shows the average across all NAM lines with the specific percentages listed to the right of them.

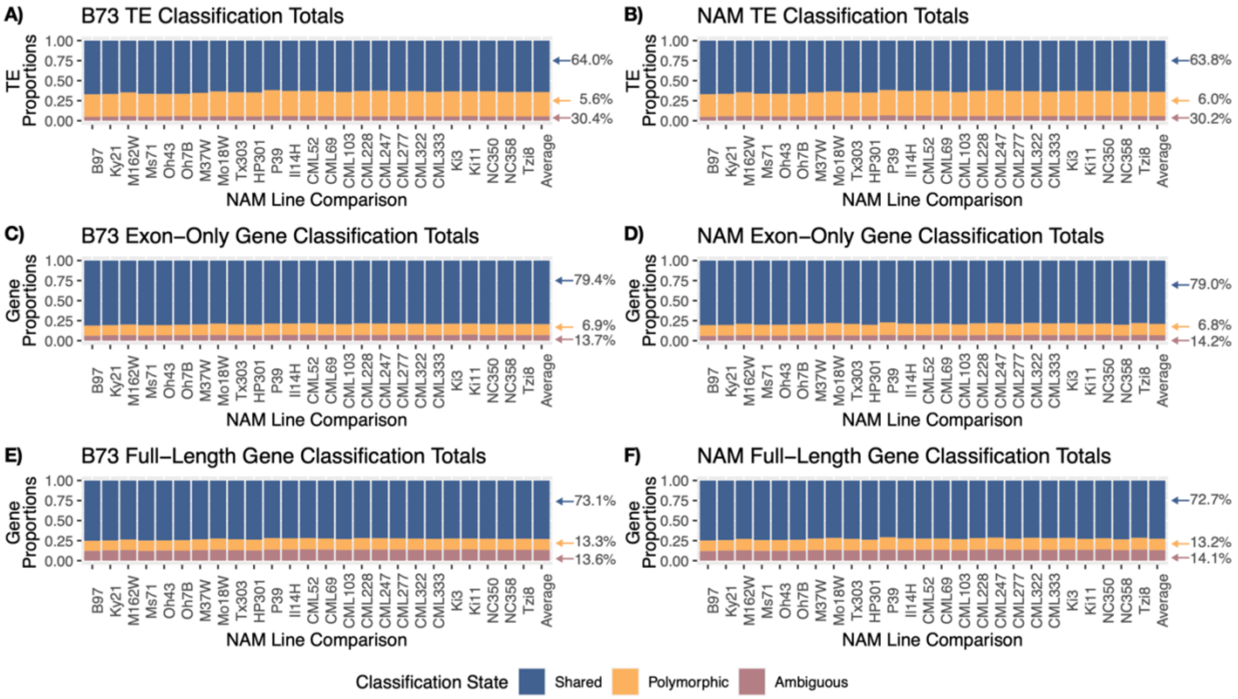


**Figure S2 Proportions of Feature Classifications Across All Pairwise Comparisons.** TE annotations (A,B), exon-only gene annotations (C,D), and full-length gene annotations (E,F) were intersected with pairwise AnchorWave alignments to classify each feature as either shared, (light blue), polymorphic (yellow), or ambiguous (light purple). Proportions of B73 features against every NAM line (A,C,E) and proportions of every NAM feature against B73 (B,D,F) classified as shared, polymorphic, or ambiguous. The last bar shows the average across all comparisons with the specific percentages listed to the right of them.

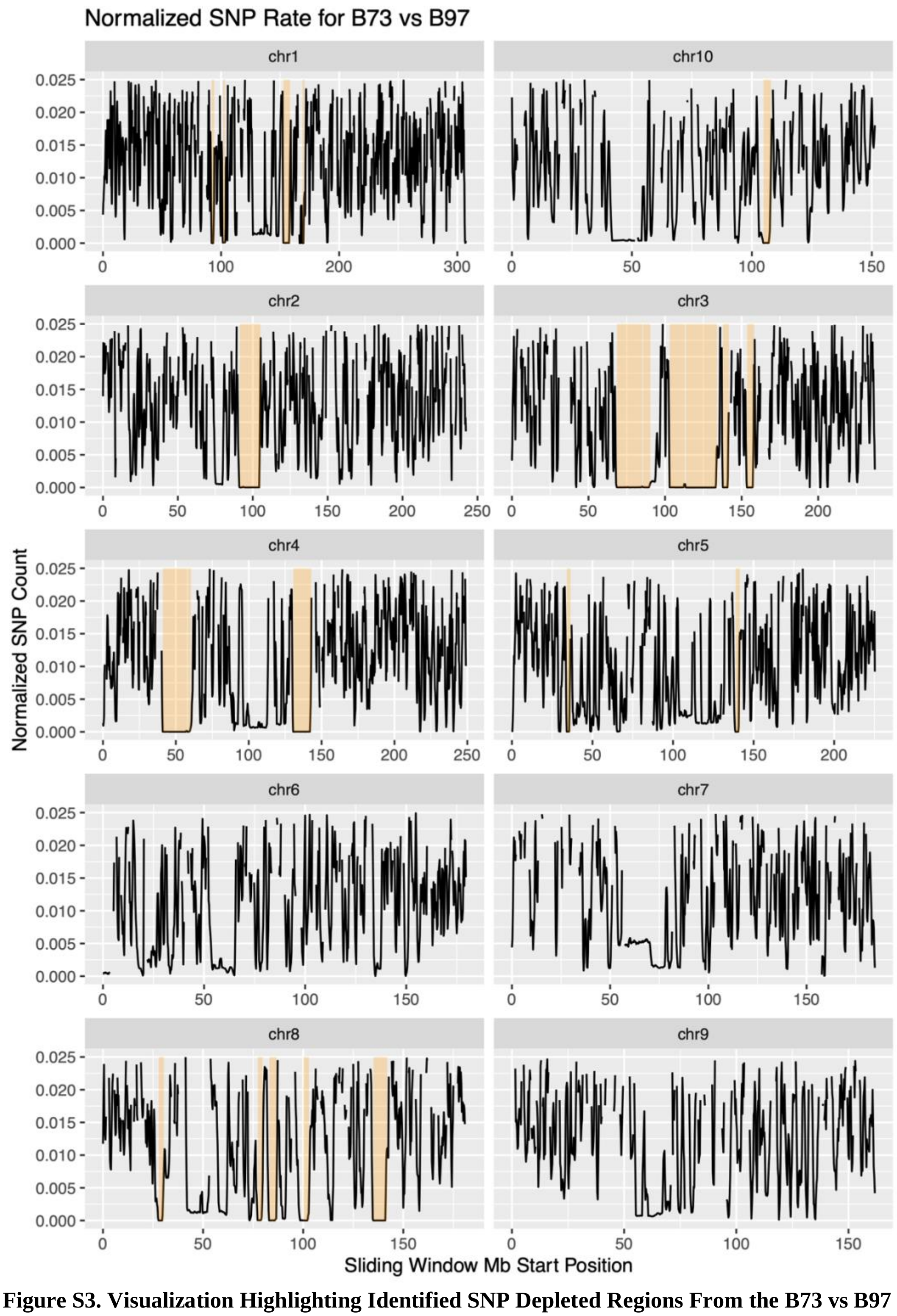


**Figure S3. Visualization Highlighting Identified SNP Depleted Regions From the B73 vs B97 AnchorWave Pairwise Alignments.** Each panel shows a different chromosome with the x-axis indicating the start position of a 1Mb window with the y-axis showing the normalized SNP count (raw SNP count/base pairs of alignable sequence). Yellow boxes indicate regions identified as SNP depleted exhibiting a normalized SNP rate less than 1 in 10,000. Gaps represent windows of either missing data or regions where there was no alignable sequence.

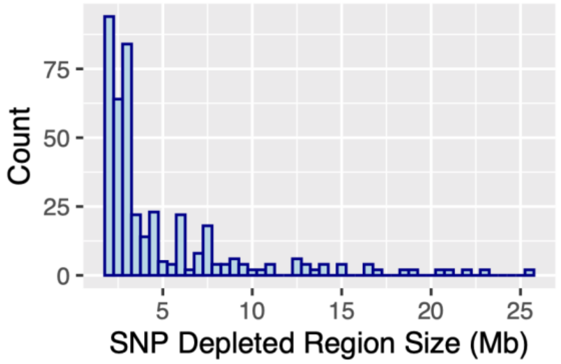


**Figure S4. Distribution of Sizes for SNP Depleted Regions Across AnchorWave Pairwise Alignments.** The x-axis shows the size of the region in megabases, while they y-axis shows the count

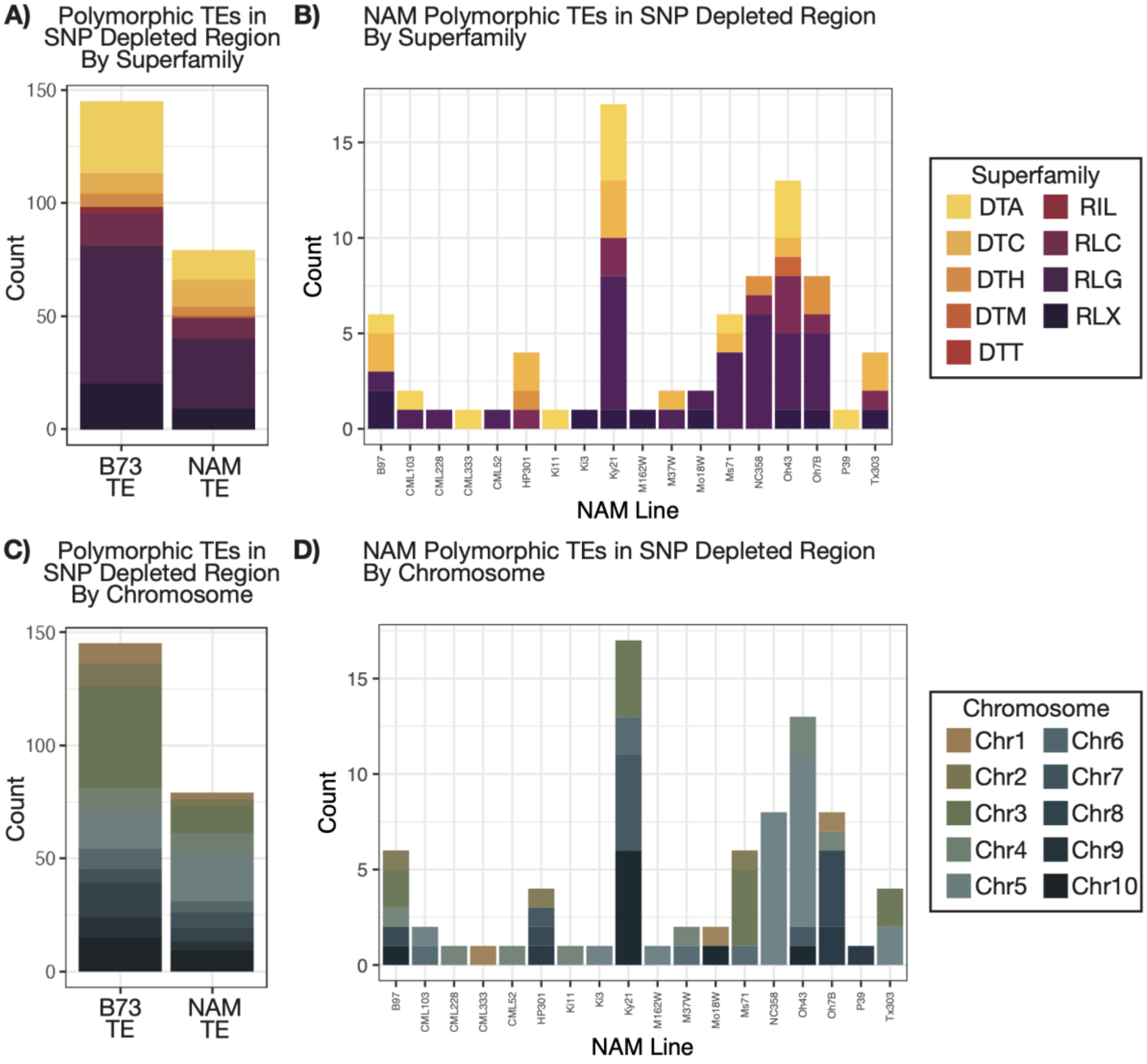


**Figure S5. Distribution of Polymorphic TEs in SNP Depleted Regions.** The first row shows this distribution by superfamily, while the second row shows this distribution by chromosome. (A) and (C) partition polymorphic TEs based on whether they are present in B73 or if they are present in a NAM line. For polymorphic TEs in NAM lines, (B) and (D) expand on this to show the distribution across NAM lines.

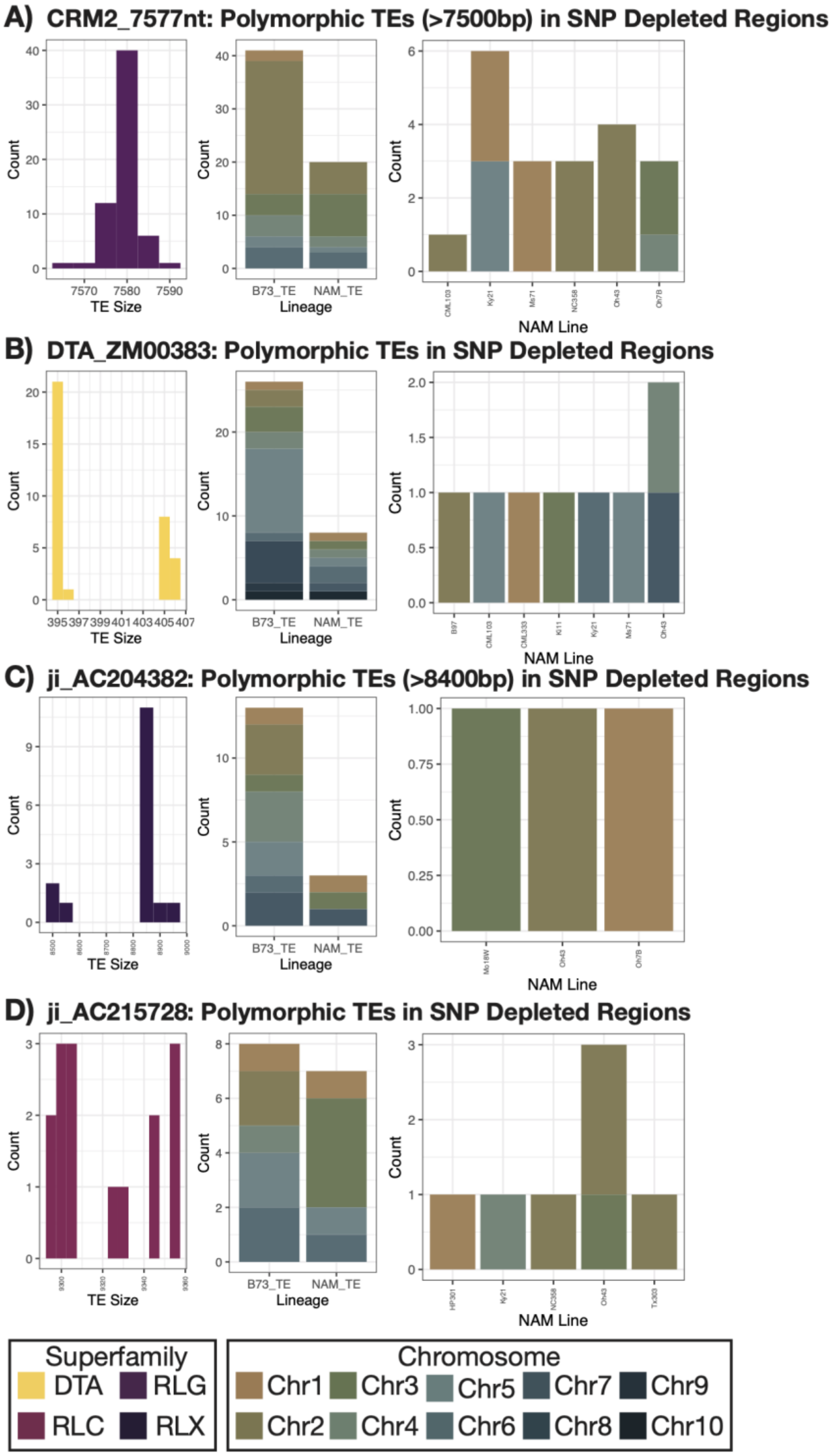


**Figure S5. Distribution of Polymorphic TEs in SNP Depleted Regions for Potentially Active Families.** Four families were identified as potentially being active due to a high number of polymorphic TEs in SNP depleted regions: (A) CRM2_7577nt (N=67), (B) DTA_ZM00383 (N=34), (C) ji_AC204382 (N=17), and (D) ji_AC215728 (N=15) . Some TEs were dropped to improve visualization including 6/67 polymorphic TEs for CRM2_7577nt and 1/17 polymorphic TEs for ji_AC204382. Column 1 in each plot shows the size distribution of polymorphic TEs in SNP depleted regions. Column 2 shows the number of polymorphic TEs in either B73 or NAM partitioned by chromosome. For polymorphic TEs present in NAM, column 3 shows the specific distribution across lines.
